# Optimizing multiplex CRISPR/Cas9-based genome editing for wheat

**DOI:** 10.1101/051342

**Authors:** Wei Wang, Alina Akhunova, Shiaoman Chao, Eduard Akhunov

## Abstract

**Background:** CRISPR/Cas9-based genome editing holds great promise to accelerate the development of new crop varieties by providing a powerful tool to modify the genomic regions controlling major agronomic traits. To diversify the set of tools available for wheat genome engineering, we have established a tRNA-based multiplex gene editing strategy for hexaploid wheat.

**Results:** The functionality of the various CRISPR/Cas9 components was assessed using the transient expression in the wheat protoplasts followed by next-generation sequencing (NGS) of the targeted genomic regions. The efficiency of wheat codon-optimized Cas9 for targeted gene editing in wheat was validated. Multiple single guide RNAs (gRNAs) were evaluated for the ability to edit the homoeologous copies of four genes affecting some important agronomic traits in wheat. Low correspondence was found between the gRNA efficiency predicted bioinformatically and that assessed in the transient expression assay. A multiplex gene editing construct with several gRNA-tRNA units under the control of a single promoter for the RNA polymerase III generated indels at the targets sites with the efficiency comparable to that obtained for a single gRNA construct.

**Conclusions:** By integrating the protoplast transformation assay with multiplexed NGS, it is possible to perform fast functional screens for a large number of gRNAs and to optimize constructs for effective editing of multiple independent targets in the wheat genome. The multiplexing capacity of the tandemly arrayed tRNA–gRNA construct is well suited for the simultaneous editing of the redundant gene copies in the allopolyploid genomes or genomic regions beneficially affecting multiple agronomic traits. A polycistronic gene construct that can be quickly assembled using the Golden Gate reaction along with the wheat codon optimized Cas9 will further expand the set of tools available for engineering the wheat genome.

## Background

Genome editing approaches have been revolutionized by the discovery of <u>c</u>lustered <u>r</u>egularly <u>i</u>nterspaced <u>s</u>hort <u>p</u>alindromic <u>r</u>epeats (CRISPR) that can guide the Cas9 nuclease to specific targets in the genome and create double strand breaks (DSB) [1]. The DSB can be repaired through either the error-prone non-homologous end joining (NHEJ) process generating loss-of-function deletions or, if the exogenous template complementary to the DSB flanking sequences is provided, the homology-directed repair (HDR) resulting in the insertion or replacement of a specific sequence. This system provides a very simple, highly specific, versatile tool to introduce targeted changes to the genomes of any species amenable to transformation.

Since the first introduction of the CRISPR/Cas9 system for editing mammalian genomes [2, 3], it has been applied to many model or crop plants including tobacco [4, 5], tomato [6, 7], barley [8], *Arabidopsis* [4], wheat [9–11], rice [11, 12] and maize [13]. In these studies, some components of the CRISPR/Cas9 editing system have been optimized for plants to improve the specificity, throughput, and efficiency. Even though a substantial increase in the expression level of Cas9 and the efficiency of genome editing have been achieved by introducing a plant codon optimized Cas9 [4], most studies have successfully used heterologous Cas9 systems that were not optimized for the species of interest. For example, the rice codon-optimized Cas9 was applied to target the *Mlo* gene in wheat [9], and the *inox* and *pds* genes in wheat were mutagenized using the human-optimized SpCas9 [10].

The synthetic single guide RNA (gRNA) includes a 20 nt-long spacer sequence, which defines the Cas9/gRNA complex target specificity, and a scaffold sequence required for Cas9 binding [14]. While the minimal requirement for a gRNA target site is the presence of protospacer adjacent motif (PAM) NGG [15], there are other sequence features that can affect the efficiency of genome editing. It was shown that the GC content of gRNA can impact the Cas9 cleavage efficiency [16, 17], as well as, base C at position 16 [17] and base G at position 20 were preferentially found in the functional gRNAs [16–19]. Similarly, the frequency of targeted mutagenesis in *C. elegans* was increased by the presence of a GG motif at the 3’-end of gRNA [20]. Therefore, to improve gRNA design, a large scale analyses of the functional gRNAs’ sequence features were performed to develop bioinformatics algorithms for selecting highly efficient gRNA spacers [17, 19, 21–28]. By taking into account such parameters as DNA sequence composition, thermodynamic stability, the presence of duplicated targets and the accessibility of genomic regions, these tools were shown to significantly improve the gRNA’s targeting capability and specificity [19, 21, 22]. Only a few of these tools were optimized to design gRNAs for wheat [25, 29] or other plants [30].

To increase the number of targets that can be simultaneously edited, several gRNA multiplexing strategies have been developed. Multiple gRNAs, each under the control of its own promoter, were combined into a single construct to edit multiple targets in maize and *Arabidopsis* [13]. Effective editing of multiple genes in rice was accomplished by putting multiple gRNAs separated by tRNA spacers under the control of a single promoter [12]. Processing of this polycistronic gene through the endogenous tRNA processing system resulted in functional gRNAs capable of guiding Cas9 to multiple targets. These multiplex gene editing strategies provide a powerful tool for implementing diverse gene editing scenarios ranging from targeting the members of multigene families [31] or gene networks to replacing or deleting multiple genomic regions simultaneously [3].

Here we have optimized the multiplex gene editing procedure for hexaploid wheat. To expand the set of tools available for CRISPR/Cas9-based editing of the wheat genome, we have developed a wheat optimized Cas9 gene and demonstrated its functionality by editing the *Q* [32], *TaGW2* [33], *TaLpx-1* [34] and *TaMLO* [9] genes. The gRNA selection procedure for multiplex gene editing included the transient expression assay in wheat protoplasts followed by deep NGS of multiplexed barcoded PCR products generated for different target regions. We demonstrate that effective multiplex gene editing in wheat can be achieved using a single polycistronic gene construct including the array of gRNA-tRNA units [12] under the control of a single promoter.

## Methods

### Plasmids and vector construction

A fragment of vector pAHC17 between the cut sites for SphI and XhoI restriction enzymes (NEB) containing the maize ubiquitin gene promoter (*ubi*) was used to replace the 35S promoter region in vector pA7-GFP (Katrin Czempinski, Potsdam University, Germany). This new plasmid construct is referred to as pA9-GFP (Figure 1A). Wheat codon optimized Cas9 coding sequence (CDS) fused from both sides with the nuclear localization signals (NLS) was synthesized by Integrated DNA Technology (IDT). The XhoI and SmaI cut sites added to the 5’- and 3’-ends of the fragment during the synthesis were used to subclone it into the pBluescript SK (PSK) vector. Then the Cas9-containing fragment between XhoI and XbaI cut sites was excised from the PSK vector, and used to replace the GFP gene between the XhoI and XbaI sites in the pA9-GFP construct. Henceforth this construct will be referred to as pA9-Cas9 (Figure 1A and Supplementary figure 1A).

**Figure 1.**
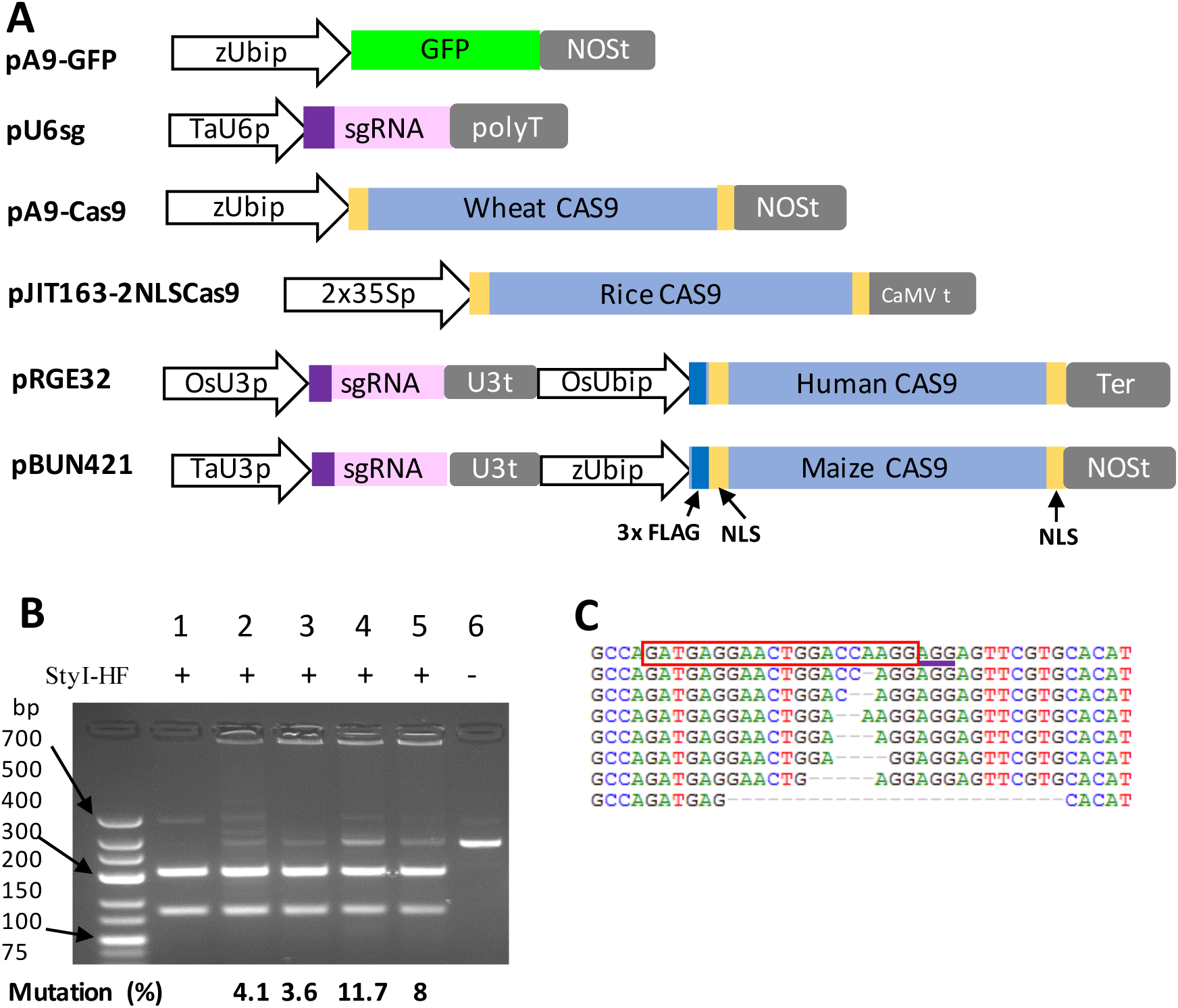
Genome editing using the wheat codon optimized Cas9. (A) Constructs used in the current study. All promoters are shown as white arrows. zUbip-maize ubiquitin gene promoter; OsUbip - rice ubiquitin gene promoter; TaU3p and TaU6p - wheat U3 and U6 promoters, respectively; OsU3p - rice U3 promoter, 2X35Sp - duplicated 35S promoter from the tobacco mosaic virus. Cas9 gene terminator sequences are shown as gray rectangles. Yellow rectangles correspond to the nuclear location signals (NLS), purple rectangles correspond to the target sequence insertion sites in the synthetic single-guide RNA (gRNA), and blue rectangles stand for 3x FLAG sequences. (B) Restriction enzyme digest of a PCR product obtained using the primers flanking the *Q* gene region targeted by Cas9 and QT1-gRNA (Supplementary Fig. 2). Lane 1 and 6: PCR products from the protoplasts transformed with pA9-GFP, Lanes 2 - 5: PCR products from the protoplasts co-transformed with the gRNA and the different versions of Cas9 from rice (Lane 2), humans (Lane 3), maize (Lane 4), and wheat (Lane 5), respectively. PCR products on Lanes 1 - 5 are digested with the StyI, Lane 6 is non-digested PCR product. Percent of mutations (%) was calculated from the band intensities (see Methods). (C) Sanger sequencing of the *Q* gene region targeted by the wheat optimized Cas9 and QT1-gRNA. The wild type sequence is shown on the top row. The target sequence is shown in the red rectangle; the PAM sequence is underlined.

The wheat U6 promoter and gRNA sequence flanked by BsaI cut sites was synthesized by IDT with the BamHI and SacI cut sites added to the 5’- and 3’- ends, respectively. The NOS poly-A terminator fragment between the SacI - EcoRI fragment was cut from the pA7-GFP construct and sub-cloned into pHSG299. The synthesized U6 promoter - gRNA fragment was cloned into vector pHSG299 between the BamHI and SacI cut sites (henceforth, pU6sg plasmid) (Figure 1A and Supplementary figure 1B).

The following plasmids were ordered from the Addgene plasmid repository: 1) pBUN421 [13], pRGE32 [12] and pJIT163-2NLSCas9 [11] with the maize, human, and rice codon optimized Cas9 genes, respectively (Figure 1A), and 2) pGTR [12] containing gRNA-tRNA fused units. The latter was modified to create a polycistronic tRNA-gRNA (*PTG*) gene construct for wheat.

**Figure 2.**
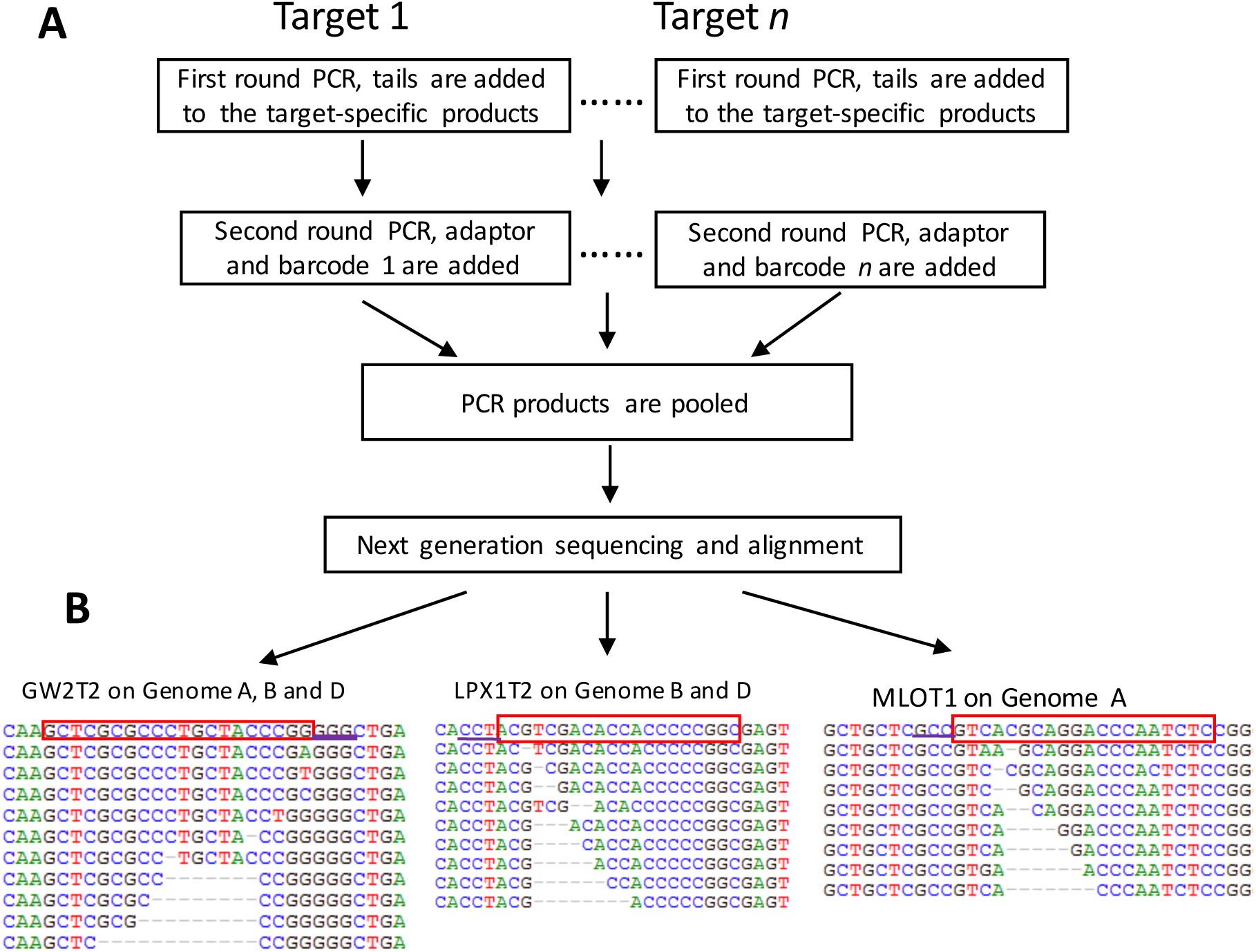
Next generation sequencing of barcoded PCR amplicons obtained for multiple Cas9-targeted genomic regions. (A) Workflow of multiplexed PCR amplicon library preparation for NGS. Multiple targets are amplified and barcoded in two rounds of PCR, pooled, and sequenced on the Illumina MiSeq instrument. (B) Illumina reads aligned to the wild type reference sequences. The target sequences are shown in the red rectangles; the PAM sequences are underlined.

The targeting portion of the single gRNAs was synthesized as a pair of reverse complementary oligonucleotides (Supplementary Table 1 and Supplementary Fig. 1B), which created the 5’ overhang tails after annealing. The reaction mix contained 2 μL of 10× annealing buffer (200 mM Tris-HCl, 100 mM (NH4)2SO4, 100 mM KCl, 1% Triton X-100, 20Mm MgCl2, pH9.0), 2 μL of 100μM forward and reverse oligos, and 14 μL ddH2O. The mix was denatured for 5 min at 94 ºC followed by the gradual 1ºC per min temperature decrease until it reached 30 ºC. Annealed oligos were sub-cloned into pU6sg or pBUN421 using the Golden Gate cloning method [35]. The Golden Gate reaction was assembled by mixing 2 μL of annealed target, 0.5 μL of pU6sg (0.2 μg/μL) or pBUN421 (0.6 μg/μL), 2 μL of 10× T4 DNA ligase buffer, 2μL of BSA (1mg/mL), 1 μL of T4 ligase (400 U/μL), 0.5 μL of BsaI-HF (20 U/μL), and 12 μL ddH2O. The mix was incubated for 50 cycles at 37 ºC for 5 min and 16 ºC for 10 min, followed by final incubation at 50 ºC for 30 min. Golden Gate reaction products were transformed into the chemically competent cells of *E. coli* strain DH5α by heat shock. Plasmid DNA was extracted using QIAprep Spin Miniprep Kit (Qiagen) and validated by Sanger sequencing.

**Table 1.** Efficiency of gRNA design for wheat genome editing. a – WU scores (0-100) are calculated using the WU-CRISPR tool [22] with a bioinformatics algorithm trained by reanalyzing empirical data from Doench *et.al*. [17]. b – E scores (0-100) are obtained using E-CRISP [29]. c – sg scores (0-100) are the gRNA’s editing activity scores calculated using the sgRNA Scorer 1.0. This program implements a bioinformatics algorithm trained on empirical data [19]. d – Mul scores (0-1) are the gRNA’s editing activity scores calculated using CRISPR MultiTargeter [41]. The scores are predicted using the bioinformatics algorithm developed by Doench *et.al*. [17]. e – “0” means that a gRNA did not have genome editing events detected or that a gRNA did not meet gRNA designers’ selection criteria.

| gRNA Name | Gene | Editing event | Efficiency (NGS) | Wheat genome | WU score<sup>a</sup> | E score<sup>b</sup> | sg score<sup>c</sup> | Mul score<sup>d</sup> |
| --- | --- | --- | --- | --- | --- | --- | --- | --- |
| Q3endT | *Q* (3' end) | Yes | 4.3% | A | 0 | 0 | 67.4 | 0.04 |
| Q3endT3 | *Q* (3' end) | Yes | 4.3% | A | 0 | 48.2 | 78.5 | 0.41 |
| Q5endT | *Q* (5' end) | Yes | 18.5% | AD | 0 | 0 | 36.1 | 0.1 |
| Q5endT4 | *Q* (5' end) | Yes | 28.5% | AD | 0 | 0 | 30.7 | 0.17 |
| Q5endT5 | *Q* (5' end) | No | 0<sup>e</sup> |  | 88 | 0 | 0 | 0 |
| Q5endT6 | *Q* (5' end) | Yes | 44.5% | AD | 0 | 54 | 59.9 | 0.38 |
| GW2T2 | *TaGW2* | Yes | 2.7% | ABD | 89 | 86.8 | 96.8 | 0.83 |
| GW2T3 | *TaGW2* | No | 0 |  | 0 | 0 | 33.7 | 0.61 |
| LPX1T1 | *TaLpx-1* | Yes | 0.3% | BD | 0 | 0 | 50.5 | 0.29 |
| LPX1T2 | *TaLpx-1* | Yes | 6.1% | BD | 0 | 52 | 59.3 | 0.18 |
| MLOT1 | *TaMLO* | Yes | 4.1% | A | 0 | 60.5 | 70.8 | 0.28 |

We have designed gRNAs for editing the set of four wheat genes including *Q* [32], *TaGW2* [33], *TaLpx-1* [34], and *TaMLO* [9]. A total of seven gRNAs were designed for the *Q* gene [32], and two gRNAs were designed to target each the *TaGW2* [33, 36], and *TaLpx-1* [34] genes (Supplementary Figure 2, Table 1 and Supplementary Table 1). A gRNA target sequence for the *TaMLO* gene on the wheat A genome was selected as a control from the previously published study (Table 1 and Supplementary Table 1) [9]. All gRNAs for genes *Q* and *TaGW2* were sub-cloned into the pU6sg and co-transformed with pA9-Cas9 into the wheat protoplasts. Whereas, the gRNAs designed for the *TaLpx-1* and *TaMLO* genes were delivered into the wheat protoplasts as single-plasmid constructs sub-cloned into pBUN421.

We have modified the previously reported method for assembling the *PTG* gene construct [12]. Our modification allows the subcloning of multiple PCR fragments, each including a gRNA-tRNA block, into the pU6sg or pBUN421 plasmids in a single-step Golden Gate reaction. As shown in Supplementary Table 1 and Supplementary Figure 3, all primers for the PCR amplification of gRNA-tRNA blocks had 5’-end tails with the BsaI cut sites for Golden Gate assembly. The forward primer of the first gRNA-tRNA block and the reverse primer of the last gRNA-tRNA block included the complete 20-nt long target gRNA sequences. The target sequence of the middle gRNA was divided among the two PCR primers used to amplify two neighboring gRNA-tRNA blocks. These primers had four complementary nucleotides, which created sticky ends after the digestion with BsaI and were used for PCR fragment ligation in the Golden Gate reaction (Supplementary Figure 3). PCR mix included the 10 μL NEB Next High Fidelity 2× PCR Master Mix, 2 μL plasmid pGTR (5 ng/μL), 2 μL each primer (5 μM), and 4 μL ddH2O. PCR conditions were as follows: 98 ºC for 30 sec followed by 35 cycles of 98 ºC for 10 sec, 56 ºC for 15 sec, 72 ºC for 10 sec, and final extension step at 72 ºC for 5 min. The resulting PCR products were sub-cloned into pU6sg or pBUN421 in a single-step Golden Gate reaction by mixing each PCR product in equal amounts (20 – 50 ng) with 0.5 μL of pU6sg (0.2 μg/μL) or pBUN421 (0.6 μg/μL), 2 μL of 10× T4 DNA ligase buffer, 2 uL of BSA (1mg/mL), 1 μL of T4 ligase (400 U/μL), 0.5 μL of BsaI-HF (20 U/μL), and (14 - x) μL of ddH2O. The products of the Golden Gate reaction were transformed into the DH5α strain of *E. coli*, followed by testing the accuracy of construct assembly using Sanger sequencing. This procedure was used for assembling plasmid pBUN421-GLM with gRNAs *TaGW2T2, TaLPX1T2*, and *TaMLOT1*.

The pU6sg-tQT1sgt construct including a single gRNA targeting the *Q gene* was designed for testing the efficiency of tRNA-based gRNA processing system in wheat [12]. The pU6sg-tQT1sgt was constructed using a strategy similar to that described for pBUN421-GLM (primers are shown in Supplementary Table 1).

### Protoplast transformation and DNA isolation

Protoplast transformation was performed using a modified protocol developed for *Arabidopsis* [37]. The seedlings of *Triticum aestivum* cultivar Bobwhite were grown in dark at 25°C for 1-2 weeks. Shoot tissues from 50 seedlings were sliced into 1-mm strips with a razor blade and placed into a flask with the digestion solution including 0.15 M Sorbitol, 0.25 M sucrose, 35 mM CaCl2, 20 mM KCl, 1.5% Cellulase R10 (From Trichoderma Viride, 7.5 U/mg), 0.75 % Macerozyme (R10 Macerating enzyme from Rhizopus sp. RPI) and 10 mM MES-KOH (pH 5.7). All chemicals were ordered from Sigma Aldrich. To infiltrate the digestion solution into leaf tissues, a vacuum (-600 mbar) was applied to the flask for 30 min in the dark followed by incubation at room temperature for 2 hours with gentle shaking at 20-30 rpm. The digested tissues were filtered through a 40 um nylon mesh into a centrifuge tube followed by rinsing the mesh with 20 mL of W5 solution (0.1 % glucose, 0.08 % KCl, 0.9 % NaCl, 1.84 % CaCl_2_•2H_2_O, 2 mM MES-KOH, pH 5.7). Due to the difference in the concentration of sucrose in the tube, the digestion solution (0.25 M sucrose) and W5 are separated into two phases. The tube was centrifuged for 7 min at 100 g at the room temperature. The protoplast fraction accumulated at the interface of the digestion solution and W5 was collected. It was then diluted 2 times in W5 and precipitated by centrifuging for 5 min at 60 g. Protoplasts were resuspended in 10 mL of W5 and precipitated again by centrifugation. Finally, protoplasts were resuspended in 3 ml of W5 and incubated on ice for 30 min.

Before transformation, W5 was removed and the protoplasts were resuspended in the MMG solution (0.4 M mannitol, 15 mM MgCl_2_, 4 mM MES-KOH, pH 5.7). The protoplast count was adjusted to 10^6^ protoplasts /mL. Ten micrograms of plasmid DNA in 10-20 μL volume was gently mixed with 100 μL of protoplasts followed by adding 130 μL of PEG-calcium transfection solution (40 % PEG4000, 0.2 M mannitol, 100 mM CaCl_2_). The resulting solution was gently mixed by agitating a tube and incubated at room temperature for 30 min. Then the transfection mix was diluted with 500 μL of W5, the protoplasts were collected by centrifuging at 100 g for 2 min, and then resuspended in 1 mL of W5.

After 18 hours of incubation at room temperature, the GFP signal in the protoplasts transformed with pA9-GFP was estimated using the Zeiss LSM 780 confocal microscope with the excitation wave length of 488 nm. The efficiency of transformation calculated as the fraction of GFP expressing protoplasts was close to 60%.

Protoplasts were incubated for 48 hours before DNA isolation. For DNA extraction, protoplasts were collected by centrifugation, mixed with 300 μL of TPS (100 mM Tris-HCl, 10 mM EDTA, 1 M KCl, pH8.0) by vortexing followed by adding 300 μL of isopropanol. After 20 min incubation, DNA was precipitated by centrifugation for 10 min at 13,000 rpm, rinsed with 1 mL of 70% ethanol, dried at room temperature, and resuspended in 20μL ddH2O.

### Validation of genome editing events

To detect mutations induced by the CRISPR/Cas9 system, genomic regions harboring the gRNA targets were PCR amplified and sequenced (Figure 1C). PCR was performed in 20 μL reaction volume using 10 μL NEB Next High Fidelity 2× PCR Master Mix, 2 μL protoplast DNA and 2 μL of each primer (5 μM) and 4 μL ddH2O.

The efficiency of target editing using gRNA QT1 was assessed by digesting PCR products with the StyI (NEB) enzyme (Figure 1B). The digestion was performed in a 20 μL reaction volume by mixing 5 μL of PCR product, 2 μL of 10× Cut Smart Buffer, 0.5 μL of StyI-HF (20 U/μL), and 12.5 μL of ddH2O. The reaction mix was incubated at 37 ºC overnight, and analyzed on a 3% agarose gel. The band intensities on the agarose gel images were measured with background subtracted using GelQuant.NET (biochemlabsolutions.com). To calculate the frequency of genome editing, the non-digested band intensity was divided by the sum of all band intensities.

### Estimating gene editing efficiency by NGS

To simultaneously analyze the large number of edited genomic regions using the NGS approach, amplicons from the targeted regions were barcoded during the two rounds of PCR with the Illumina’s TruSeq adaptors. During the first round, we have used target-specific forward and reverse PCR primers modified by adding to the 5’-end a 30-nucleotide long sequence complimentary to parts of the Illumina’s TruSeq adaptors (Supplementary Table 1). The second round of PCR was performed using the primers complimentary to the 5’-ends of the first set of primers. This second set of primers completes the reconstruction of TruSeq adaptor sequences and adds the amplicon-specific barcodes (Supplementary Fig. 4). PCR products were purified with MinElute PCR Purification Kit (Qiagen), normalized to 10 nM concentration, pooled in equal volumes, and then sequenced at the K-State Integrated Genomics Facility on the MiSeq instrument using the 600 cycles MiSeq reagent v3 kit.

De-multiplexed and quality-trimmed reads were aligned to each of the three homoeologous copies genes using BWA [38] followed by SNP and indel calling with SAMtools [39, 40]. The efficiency of target editing was estimated as the proportion of reads with deletions relative to the total number of aligned reads using a custom Perl script.

## Results

### Genome editing using the wheat codon optimized Cas9

The pU6sg-QT1 construct expressing gRNA targeting the *Q* gene (Simons et al., 2006) was co-transformed into wheat protoplasts with the constructs expressing wheat (pA9-Cas9), maize (pBUN421) [13], human (pRGE32) [12] or rice (pJIT163-2NLSCas9) [11] codon-optimized Cas9. The ability of all tested Cas9 versions to perform gRNA-guided genome editing was confirmed by the StyI digestion of amplicons from the target site (Figure 1B). Based on the gel image band intensities, the efficiency of construct pA9-Cas9 (8%) with wheat-optimized Cas9 was lower than that of pBUN421 (11.7%), but was higher than the efficiency of constructs pRGE32 (3.6%) or pJIT163-2NLSCas9 (4.1%). The Sanger sequencing of the subcloned non-digested PCR products from the pA9-Cas9 transformed protoplast DNA revealed a number of deletions resulting from NHEJ-mediated reparation of DNA (Figure 1C).

### Analyzing genome editing events by next-generation sequencing (NGS)

The transient expression in the transformed protoplasts provides a simple and rapid method for assessing the editing capability of the CAS9/gRNA constructs [10, 11]. Here we have adopted this approach to screen gRNAs for the ability to generate Cas9-mediated changes in the wheat genome. Further, the protoplast expression assay was combined with NGS for the rapid and cost-effective analysis of multiple genomic regions (Figure 2A and Supplementary Figure 4). This strategy was used to evaluate the gRNAs designed to target four genes controlling domestication (*Q* gene) [32], seed development (*TaGW2*) [33] and disease resistance (*TaLpx-1* and *TaMLO*) phenotypes in wheat [9, 34].

The editing specificity of the designed gRNAs was consistent with the divergence between the homoeologous gene copies at the target site regions (Table 1). The A and D genome copies of the *Q* gene were successfully targeted by three gRNAs on the 5’-end (Figure 2B and Supplementary Figure 5 and 6), and only the A genome copy was targeted by two gRNAs on the 3’-end (Supplementary Figure 7). Only one of the two target sites in *TaGW2* was edited in all three wheat genomes (Figure 2B and Supplementary Figure 8). Both gRNAs designed for the *TaLpx-1* gene produced deletions in the B and D genomes, whereas no editing events were detected for the target in the A genome due to its high divergence from the guide sequence (Figure 2B and Supplementary Figure 9). The gRNA targeting *TaMLO* induced mutations only in the wheat A genome, as previously reported (Figure 2B and Supplementary Figure 10) [9].

### Validating the gRNA design tools

To test whether the failure of some gRNAs to produce deletions is caused by suboptimal gRNA design, we have used algorithms implemented in WU-CRISPR [22], E-CRISP [29], sgRNA Scorer 1.0 [19], and CRISPR MultiTargeter [41], to analyze the gRNAs used in our study. Among these programs only E-CRISP has a feature for selecting gRNAs to target the wheat genome.

Out of eleven gRNAs analyzed in the protoplast transformation assay, only two had prediction scores assigned by WU-CRISPR, five gRNAs matched the selection criteria of E-CRISP, and ten gRNAs had prediction scores assigned by sgRNA Scorer 1.0 or CRISPR MultiTargeter respectively (Table 1). While only one out of two gRNAs selected by WU-CRISPR produced mutations in a target gene, all five gRNAs designed by E-CRISP resulted in genome editing events. Among those gRNAs that failed to meet the WU-CRISPR’s and E-CRISP’s design criteria, eight and four, respectively, were shown to be mutagenic in the protoplast assays. Out of the ten gRNAs designed by sgRNA Scorer 1.0 and CRISPR MultiTargeter, nine were validated in the protoplast assays. However, in spite of the high validation rate for these programs, the gRNA efficiency estimated by NGS and the gRNA scores (Table 1) showed no correlation for sgRNA Scorer 1.0 and CRISPR MultiTargeter (*R*^2^ = −0.06 and −0.11, respectively).

### Multiplex gene editing in hexaploid wheat

There are several approaches developed for multiplex gene editing in humans, maize, *Arabidopsis*, and rice [3, 4, 12, 13]. The recently developed method based on a single polycistronic gene comprised of multiple tandemly arrayed tRNA – gRNA blocks compatible with the endogenous tRNA processing system provides one of the most efficient and versatile tools for multiplex gene editing [12]. Here we report the optimization of the PTG/Cas9 system for multiplex genome editing in hexaploid wheat.

To test the ability of the gRNA subjected to tRNA processing in hexaploid wheat to serve as guides for Cas9, we used a construct (pU6sg-tQT1sgt) with the QT1 gRNA (Supplementary Table 1) flanked by the tRNA units (Figure 3A). The results of NGS confirmed successful editing of the targeted region in the *Q* gene (Figure 3B) and indicated that the endogeneous tRNA processing system of wheat can produce functional gRNAs from the expressed tRNA – gRNA units.

**Figure 3.**
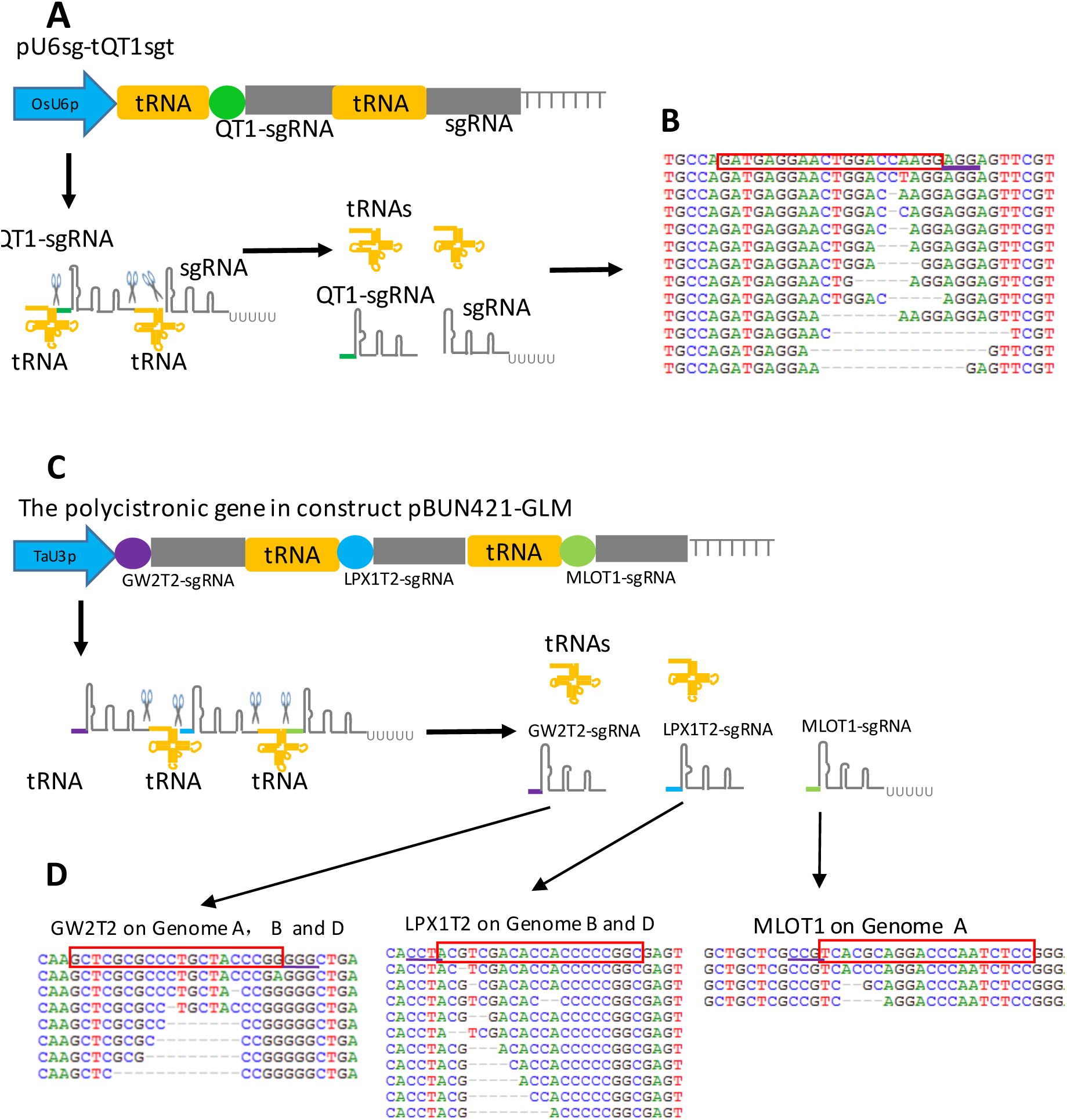
CRISPR/Cas9-mediated multiplex editing in hexaploid wheat using the endogenous tRNA-processing system. (A) The structure of a test construct including the QT1-gRNA (targets the *Q* gene) flanked by two tRNAs is shown. TaU6p - wheat U6 promoter; QT1 is shown as a green circle; gRNA is shown as a grey rectangle. The gRNAs are released after the tRNA processing of a transcript encoded by a polycistronic gene including the QT1-gRNA and one gRNA without a guide sequence. (B) Alignment of Illumina reads obtained for the QT1-gRNA-targeted regions of the *Q* gene. The wild type sequence is shown on the top. The QT1 target sequence is shown in the red rectangle; the PAM sequence is underlined. (C) Schematic of tRNA-based processing of a polycistronic gene construct (pBUN421-GLM) containing three tRNA-gRNA blocks. Target sequences GW2T2, LPX1T2, and MLOT1 are shown in purple, blue, and green color, respectively. (D) The NGS of three genomic regions targeted by the pBUN421-GLM construct. The wild type sequence is shown on the top. The target sequences are shown in the red rectangles; the PAM sequences are underlined.

A higher level of multiplexing was tested by combining the GW2T2-gRNA, LPX1T2-gRNA and MLOT1-gRNA interspaced by tRNAs into a polycistronic gene construct pBUN421-GLM (Figure 3C). For the comparison of editing efficiency, single-gRNA construct pBUN421-LPX1T2 was transformed into protoplasts. The NGS results obtained for the pBUN421-GLM transformants (Figure 3D, Supplementary figure 11 and 12) indicated that the polycistronic transcript was correctly processed resulting in functional gRNAs capable of directing Cas9 to the respective genomic targets. The editing efficiencies obtained by multiplex editing for the *TaGW2*, *TaLpx-1* and *TaMLO* gene targets were 7.4%, 4%, and 0.5%, respectively. In the same transformation experiment, the efficiency of *TaLpx-1* editing using the LPX1T2 gRNA as a part of the multi-gRNA editing construct (4%) or as a single-gRNA editing construct (3.9%) were comparable.

## Discussion

Because the development of transgenic wheat lines is a time-consuming and laborious process, the selection of functional gRNAs before plant transformation, especially for multiple targets, is critical for successful genome editing. The protoplast transient expression assay in combination with the multiplexed NGS of pooled amplicons was shown to be an effective strategy for large-scale gRNA pre-screening. Several studies reported the usage of NGS for evaluating the efficiency of Cas9-based editing. The Nextera kit (Illumina) was used for the NGS of PCR-amplified regions targeted by the gRNA/Cas9 complex [42]. A high-throughput strategy for detecting the CRISPR-Cas9 induced mutations in mouse used unique barcodes added to the PCR products of different target sites followed by ligating Illumina adaptors and NGS [43]. The approach utilized in our study allows for adding barcodes and Illumina adaptors during the two rounds of PCR and generating products that can be sequenced directly avoiding the time-consuming library preparation step. Another advantage of the NGS-based analysis of the Cas9 editing events in allopolyploid wheat is that it provides valuable information about the efficiency of homoeologous target editing. Using sequence data, it is possible to infer which homoeolog has undergone changes and optimize the gRNA design to modify either a specific homoeologous copy of a gene or all homoeologs simultaneously. The throughput of the gRNA screening procedure can be increased by increasing the number of barcoded primers in the second round of PCR to include a larger number of targets as needed.

While the majority of gRNAs matching the E-CRISP, sgRNA Scorer 1.0, or CRISPR MultiTargeter design criteria were able to produce editing events in the wheat genome, we found some discrepancy between the bioinformatically predicted gRNA scores and the editing efficiency in the protoplast transformation assays. This result could partially be explained by the usage of empirical data obtained in humans [17, 19] for optimizing these bioinformatics algorithms. In addition, the effective gRNA design using E-CRISP, that has wheat-specific design functionality, was achieved at the cost of increased false negative rate; many gRNAs classified as non-functional by this tool showed good editing capability in the wheat protoplasts. This discrepancy could potentially be associated with the E-CRISP’s algorithm that during the calculation of the gRNA activity score applies a penalty if duplicated targets that are identical or nearly identical to the original target exit in the genome [29]. Due to the polyploid nature of the wheat genome this criterion may reduce the number of gRNAs designed to the duplicated genomic regions [29]. It appears that further improvements of the bioinformatical tools will be required for designing gRNAs optimized for genome editing in polyploid wheat.

Here we demonstrated that the editing of multiple gene targets in the wheat genome could be achieved using a construct with an array of the gRNA-tRNA units expressed from the wheat U3 snoRNA promoter. It was demonstrated that a transcript produced from this construct can generate functional gRNAs after being processed through the endogenous tRNA-processing system of wheat, consistent with the results reported for rice [12]. The high multiplexing capacity of the PTG/Cas9 system makes it well suited for studying the polyploid wheat genome function. Due to the presence of duplicated gene copies, editing in the wheat genome might require multiple gRNAs for knocking out or deleting a specific set of homoeologs. Moreover, by designing gRNAs to sequences carrying homoeolog-specific nucleotide changes or to sequences conserved across all the wheat genomes, it will be possible to selectively edit either multiple genes from a specific wheat genome or to introduce modifications to all gene copies. When the useful gene variants for several agronomic traits are identified, the multiplexing capability of the PTG/Cas9 system can be used to simultaneously create beneficial allelic changes in the wheat genome in a single transformation.

## Conclusions

Our results suggest that for designing functional gRNAs for wheat genome editing, bioinformatics algorithms should take into account the polyploid nature of its genome. Currently, the transient expression assay provides an effective alternative for screening a large number of gRNAs and selecting optimal constructs for editing multiple targets in the wheat genome. Moreover, this approach combined with NGS can help to optimize gRNAs for the selective targeting of gene copies on different wheat genomes. Our study demonstrated that the tRNA processing system of wheat can generate mutagenic gRNA molecules from a polycistronic gene construct assembled using multiple gRNAs separated by the tRNA spacers. The compatibility of construct assembly process with the Golden Gate cloning system creates a flexible tool for designing sophisticated gene editing experiments. This tool is especially suited for studying gene function and modifying genomic regions controlling important agronomic phenotypes in the highly redundant allopolyploid wheat genome.

## Declarations

### Availability of data and material

The sequence data generated for CAS9-edited targets is deposited to the NCBI SRA database (http://www.ncbi.nlm.nih.gov/sra). The datasets supporting the conclusions of this article are included within the article and its additional files.

### Authors’ contributions

WW – constructed plasmids, assessed gRNA design efficiency, analyzed data and participated in drafting the manuscript; AA – designed and implemented a strategy for the NGS-based assessment of target editing efficiency, bioinfromatical analysis and participated in editing the manuscript; SC – validated construct assembly accuracy by Sanger sequencing and participated in editing the manuscript; EA – conceived idea, analyzed data and wrote the manuscript.

## Acknowledgements

We would like to thank Shichen Wang for assistance with the bioinformatical data processing and Kathrine Jordan for the valuable discussions of the presented work. We would like to thank Qi-Jun Chen for providing plasmid pBUN421, Caixia Gao for providing plasmid pJIT163-2NLSCas9, and Yinong Yang for providing plasmid pRGE32 and pGTR.

## References

1. Shalem O, Sanjana NE, Zhang F. High-throughput functional genomics using CRISPR-Cas9. Nat Rev Genet. 2015; 16:299–311.

2. Mali P, Yang L, Esvelt KM, Aach J, Guell M, DiCarlo JE, Norville JE, Church GM. RNA-guided human genome engineering via Cas9. Science. 2013; 339:823–6.

3. Cong L, Ran FA, Cox D, Lin S, Barretto R, Habib N, Hsu PD, Wu X, Jiang W, Marraffini LA et al.Multiplex genome engineering using CRISPR/Cas systems. Science. 2013; 339:819–23.

4. Li JF, Norville JE, Aach J, McCormack M, Zhang DD, Bush J, Church GM, Sheen J. Multiplex and homologous recombination-mediated genome editing in Arabidopsis and Nicotiana benthamiana using guide RNA and Cas9. Nat Biotechnol. 2013; 31:688–91.

5. Nekrasov V, Staskawicz B, Weigel D, Jones JD, Kamoun S. Targeted mutagenesis in the model plant Nicotiana benthamiana using Cas9 RNA-guided endonuclease. Nat Biotechnol. 2013; 31:691–3.

6. Brooks C, Nekrasov V, Lippman ZB, Van Eck J. Efficient Gene Editing in Tomato in the First Generation Using the Clustered Regularly Interspaced Short Palindromic Repeats/CRISPR-Associated9 System. Plant Physiol. 2014; 166:1292–7.

7. Cermak T, Baltes NJ, Cegan R, Zhang Y, Voytas DF. High-frequency, precise modification of the tomato genome. Genome Biol. 2015; 16:232.

8. Lawrenson T, Shorinola O, Stacey N, Li CD, Ostergaard L, Patron N, Uauy C, Harwood W. Induction of targeted, heritable mutations in barley and Brassica oleracea using RNA-guided Cas9 nuclease. Genome Biol. 2015; 16:258.

9. Wang Y, Cheng X, Shan Q, Zhang Y, Liu J, Gao C, Qiu JL. Simultaneous editing of three homoeoalleles in hexaploid bread wheat confers heritable resistance to powdery mildew. Nat Biotechnol. 2014; 32:947–51.

10. Upadhyay SK, Kumar J, Alok A, Tuli R. RNA-guided genome editing for target gene mutations in wheat. G3 (Bethesda). 2013; 3:2233–8.

11. Shan Q, Wang Y, Li J, Zhang Y, Chen K, Liang Z, Zhang K, Liu J, Xi JJ, Qiu JL et al.Targeted genome modification of crop plants using a CRISPR-Cas system. Nat Biotechnol. 2013; 31:686–8.

12. Xie K, Minkenberg B, Yang Y. Boosting CRISPR/Cas9 multiplex editing capability with the endogenous tRNA-processing system. Proc Natl Acad Sci U S A. 2015; 112:3570–5.

13. Xing HL, Dong L, Wang ZP, Zhang HY, Han CY, Liu B, Wang XC, Chen QJ. A CRISPR/Cas9 toolkit for multiplex genome editing in plants. BMC Plant Biol. 2014; 14:327.

14. Jinek M, Chylinski K, Fonfara I, Hauer M, Doudna JA, Charpentier E. A programmable dual-RNA-guided DNA endonuclease in adaptive bacterial immunity. Science. 2012; 337:816–21.

15. Mojica FJM, Diez-Villasenor C, Garcia-Martinez J, Almendros C. Short motif sequences determine the targets of the prokaryotic CRISPR defence system. Microbiol-Sgm. 2009; 155:733–40.

16. Wang T, Wei JJ, Sabatini DM, Lander ES. Genetic Screens in Human Cells Using the CRISPR-Cas9 System. Science. 2014; 343:80–4.

17. Doench JG, Hartenian E, Graham DB, Tothova Z, Hegde M, Smith I, Sullender M, Ebert BL, Xavier RJ, Root DE. Rational design of highly active gRNAs for CRISPR-Cas9-mediated gene inactivation. Nat Biotechnol. 2014; 32:1262–7.

18. Gagnon JA, Valen E, Thyme SB, Huang P, Akhmetova L, Pauli A, Montague TG, Zimmerman S, Richter C, Schier AF. Efficient mutagenesis by Cas9 protein-mediated oligonucleotide insertion and large-scale assessment of single-guide RNAs. PLoS One. 2014; 9:e98186.

19. Chari R, Mali P, Moosburner M, Church GM. Unraveling CRISPR-Cas9 genome engineering parameters via a library-on-library approach. Nat Methods. 2015; 12:823–6.

20. Farboud B, Meyer BJ. Dramatic Enhancement of Genome Editing by CRISPR/Cas9 Through Improved Guide RNA Design. Genetics. 2015; 199:959–71.

21. Xu H, Xiao T, Chen CH, Li W, Meyer CA, Wu Q, Wu D, Cong L, Zhang F, Liu JS et al.Sequence determinants of improved CRISPR gRNA design. Genome Res. 2015; 25:1147–57.

22. Wong N, Liu W, Wang X. WU-CRISPR: characteristics of functional guide RNAs for the CRISPR/Cas9 system. Genome Biol. 2015; 16:218.

23. Upadhyay SK, Sharma S. SSFinder: high throughput CRISPR-Cas target sites prediction tool. Biomed Res Int. 2014; 2014:742482.

24. Sander JD, Maeder ML, Reyon D, Voytas DF, Joung JK, Dobbs D. ZiFiT (Zinc Finger Targeter): an updated zinc finger engineering tool. Nucleic Acids Res. 2010; 38:W462–8.

25. Naito Y, Hino K, Bono H, Ui-Tei K. CRISPRdirect: software for designing CRISPR/Cas guide RNA with reduced off-target sites. Bioinformatics. 2015; 31:1120–3.

26. Montague TG, Cruz JM, Gagnon JA, Church GM, Valen E. CHOPCHOP: a CRISPR/Cas9 and TALEN web tool for genome editing. Nucleic Acids Res. 2014; 42:W401–7.

27. Hsu PD, Scott DA, Weinstein JA, Ran FA, Konermann S, Agarwala V, Li Y, Fine EJ, Wu X, Shalem O et al.DNA targeting specificity of RNA-guided Cas9 nucleases. Nat Biotechnol. 2013; 31:827–32.

28. Doench JG, Fusi N, Sullender M, Hegde M, Vaimberg EW, Donovan KF, Smith I, Tothova Z, Wilen C, Orchard R et al.Optimized gRNA design to maximize activity and minimize off-target effects of CRISPR-Cas9. Nat Biotechnol. 2016; 34:184–91.

29. Heigwer F, Kerr G, Boutros M. E-CRISP: fast CRISPR target site identification. Nat Methods. 2014; 11:122–3.

30. Lei Y, Lu L, Liu HY, Li S, Xing F, Chen LL. CRISPR-P: a web tool for synthetic single-guide RNA design of CRISPR-system in plants. Mol Plant. 2014; 7:1494–6.

31. Peng D, Kurup SP, Yao PY, Minning TA, Tarleton RL. CRISPR-Cas9-mediated single-gene and gene family disruption in Trypanosoma cruzi. MBio. 2015; 6:e02097–14.

32. Simons KJ, Fellers JP, Trick HN, Zhang Z, Tai YS, Gill BS, Faris JD. Molecular characterization of the major wheat domestication gene Q. Genetics. 2006; 172:547–55.

33. Su Z, Hao C, Wang L, Dong Y, Zhang X. Identification and development of a functional marker of TaGW2 associated with grain weight in bread wheat (Triticum aestivum L.). Theor Appl Genet. 2011; 122:211–23.

34. Nalam VJ, Alam S, Keereetaweep J, Venables B, Burdan D, Lee H, Trick HN, Sarowar S, Makandar R, Shah J. Facilitation of Fusarium graminearum Infection by 9-Lipoxygenases in Arabidopsis and Wheat. Mol Plant Microbe Interact. 2015; 28:1142–52.

35. Engler C, Kandzia R, Marillonnet S. A one pot, one step, precision cloning method with high throughput capability. PLoS One. 2008; 3:e3647.

36. Song XJ, Huang W, Shi M, Zhu MZ, Lin HX. A QTL for rice grain width and weight encodes a previously unknown RING-type E3 ubiquitin ligase. Nat Genet. 2007; 39:623–30.

37. Zhai Z, Sooksa-nguan T, Vatamaniuk OK. Establishing RNA interference as a reverse-genetic approach for gene functional analysis in protoplasts. Plant Physiol. 2009; 149:642–52.

38. Li H, Durbin R. Fast and accurate short read alignment with Burrows-Wheeler transform. Bioinformatics. 2009; 25:1754–60.

39. Li H. A statistical framework for SNP calling, mutation discovery, association mapping and population genetical parameter estimation from sequencing data. Bioinformatics. 2011; 27:2987–93.

40. Li H, Handsaker B, Wysoker A, Fennell T, Ruan J, Homer N, Marth G, Abecasis G, Durbin R, Genome Project Data Processing S. The Sequence Alignment/Map format and SAMtools. Bioinformatics. 2009; 25:2078–9.

41. Prykhozhij SV, Rajan V, Gaston D, Berman JN. CRISPR multitargeter: a web tool to find common and unique CRISPR single guide RNA targets in a set of similar sequences. PLoS One. 2015; 10:e0119372.

42. Ran FA, Hsu PD, Wright J, Agarwala V, Scott DA, Zhang F. Genome engineering using the CRISPR-Cas9 system. Nat Protoc. 2013; 8:2281–308.

43. Bell CC, Magor GW, Gillinder KR, Perkins AC. A high-throughput screening strategy for detecting CRISPR-Cas9 induced mutations using next-generation sequencing. BMC Genomics. 2014; 15:1002.

